# Human myelin protein P2: From crystallography to time-lapse membrane imaging and neuropathy-associated variants

**DOI:** 10.1101/2021.04.15.439958

**Authors:** Maiju Uusitalo, Martin Berg Klenow, Saara Laulumaa, Matthew P. Blakeley, Adam Cohen Simonsen, Salla Ruskamo, Petri Kursula

**Affiliations:** Faculty of Biochemistry and Molecular Medicine & Biocenter Oulu, University of Oulu, Finland; Department of Physics, Chemistry and Pharmacy, University of Southern Denmark, Odense, Denmark; European Spallation Source, Lund, Sweden; Large-Scale Structures Group, Institut Laue–Langevin, Grenoble, France; Department of Biomedicine, University of Bergen, Norway

**Keywords:** myelin protein P2, fatty acid binding protein, Charcot-Marie-Tooth disease, mutation, protein structure, lipid binding

## Abstract

Peripheral myelin protein 2 (P2) is a fatty acid-binding protein expressed in vertebrate peripheral nervous system myelin, as well as in human astrocytes. Suggested functions of P2 include membrane stacking and lipid transport. Mutations in the *PMP2* gene, encoding P2, are associated with Charcot-Marie-Tooth disease (CMT). Recent studies have revealed three novel *PMP2* mutations in CMT patient families. To shed light on the structure and function of the corresponding P2 variants, we used X-ray and neutron crystallography, small-angle X-ray scattering, circular dichroism spectroscopy, computer simulations, and lipid binding assays. The crystal and solution structures of the I50del, M114T, and V115A variants of P2 showed only minor differences to the wild-type protein, whereas the thermal stability of the disease variants was reduced. Lipid vesicle aggregation assays revealed no change in membrane stacking characteristics, while the variants showed slightly altered fatty acid binding. Time-lapse imaging of lipid bilayers indicated membrane blebbing induced by P2, which could be related to its function in stacking of two curved membrane surfaces in myelin in vivo. All variants caused blebbing of membranes on similar timescales. In order to better understand the links between structure, dynamics, and function, the crystal structure of perdeuterated P2 was refined from room temperature data collected using both neutrons and X-rays, and the results were compared to molecular dynamics simulations and cryocooled crystal structures. Taken together, our data indicate similar properties of all known CMT variants of human P2; while crystal structures are nearly identical, stability and function of the disease variants are impaired compared to the wild-type protein. Our data provide new insights into the structure-function relationships and dynamics of P2 in health and disease.

## Introduction

The neuronal axons of the peripheral nervous system (PNS) are surrounded by multilamellar Schwann cell membrane protrusions, called myelin. Myelin sheaths comprise over 40 membrane lamellae providing a high-resistance and low-capacitance sheath for saltatory impulse conduction. The myelin membrane has uniquely high lipid content (~70%) and very few proteins are enriched in compact myelin, maintaining the proper insulating structure of the myelin sheath.

Human peripheral myelin protein 2 (P2), encoded by the *PMP2* gene, is a 14-kDa fatty acid-binding protein (FABP), expressed by Schwann cells in the vertebrate PNS (Chmurzyńska, 2006) and astrocytes in human central nervous system (CNS) (Kelley et al., 2018). P2 first appeared in tetrapods, but the origins of P2 can be traced to invertebrate paralogs. P2 orthologs are limited to tetrapods, although paralogs, *i.e*. other FABPs, are present in fishes and invertebrates. P2 is less conserved among mammals than other compact myelin proteins (Gould et al., 2008), which may hint at diverging importance and function for P2 between species.

According to current knowledge, the main functions of P2 are associated with lipids. P2 stacks lipid bilayers and may transport lipids, such as fatty acids or cholesterol, within myelin membranes (Majava et al., 2010; Ruskamo et al., 2020, 2014; Suresh et al., 2010; Zenker et al., 2014). P2 binds to lipid bilayers *via* two opposing faces, sticking membranes together with a constant spacing (Ruskamo et al., 2020, 2014). P2 plays a role in maintaining the glial cell lipid homeostasis (Zenker et al., 2014) and remyelination of peripheral nerves after a nerve injury (Stettner et al., 2018). A number of recent studies have highlighted a novel intriguing function of P2 in human astrocytes (Cai et al., 2015; Kelley et al., 2018), while it is absent from mouse astrocytes. Altered *PMP2* expression patterns have been observed in various cancers (Ahn et al., 2020; Cai et al., 2015; Graf et al., 2019), as well as in pathological conditions of the inner ear (Chen et al., 2021).

Point mutations in the *PMP2* gene have been linked to Charcot-Marie-Tooth disease (CMT) (Gonzaga-Jauregui et al., 2015; Hong et al., 2016; Motley et al., 2016; Punetha et al., 2018), a genetically heterogeneous group of motor and sensory neuropathies caused by mutations in >100 target genes. Globally, CMT is the most common inherited neuropathy with a prevalence of 1:2500 (Morena et al., 2019). CMT can be classified into three main types: demyelinating (CMT1), axonal (CMT2), and intermediate (I-CMT) (Stavrou et al., 2021). 40-50% of all CMT patients have CMT1 (Pareyson et al., 2017), which is characterized by the loss of myelin and reduction of the nerve conduction velocities (NCVs) to <35 m/s. Generally, individuals with myelin-damaging mutations develop symptoms at the age of 5-25. Symptoms include slowly progressive distal muscle weakness and atrophy, as well as sensory loss, often associated to the *pes cavus* foot deformity and bilateral foot drop (Bird, 2020).

Crystal structures of bovine (Jones et al., 1988), equine (Hunter et al., 2005), and human P2 have been determined (Laulumaa et al., 2018; Majava et al., 2010; Ruskamo et al., 2014). P2 is a small β-barrel protein folded like other FABPs, with a cap formed by two α-helices. P2 has two membrane-binding sites on opposite sides of the protein; the ends of the β-barrel are positively charged while the top of the α-helical cap is hydrophobic. Inside the β-barrel, a fatty acid is bound in the crystal structures. Sub-atomic resolution crystal structures have revealed an unexpected protonation state for one of the Arg residues coordinating the bound fatty acid (Laulumaa et al., 2018; Ruskamo et al., 2014). It is likely that the fatty acid ligand mimics lipids transported by P2, although a structural role for the ligand cannot be excluded.

Crystal structures of P2 variants have shed light on the details of P2 dynamics, membrane interactions, and portal region control (Laulumaa et al., 2018; Lehtimäki et al., 2012; Ruskamo et al., 2020). Three CMT1-linked P2 mutations have been studied previously at the molecular level (Ruskamo et al., 2017). These autosomal dominant mutations include I43N (Gonzaga-Jauregui et al., 2015; Hong et al., 2016), T51P, and I52T (Motley et al., 2016; Punetha et al., 2018). These three P2 disease variants have crystal structures similar to the wildtype protein, but their stability and biochemical binding properties are affected. T51P is the most differing variant, with more open solution structure conformations, altered membrane interactions, and reduced solubility (Ruskamo et al., 2017).

Recently, two novel CMT1-associated P2 point mutations; p.M114T (c.341T>C) and p.V115A (c.344T>C) were found in Bulgarian and German families, respectively (Palaima et al., 2019). The Bulgarian family with M114T suffered low NCVs, <15 m/s, of the motor fibres of median and ulnar nerves. In contrast, the German patients with the V115A mutation showed only mild changes in NCVs (Palaima et al., 2019). A third recently discovered P2 patient mutation, an in-frame deletion of Ile50 (I50del, c. 147-149delTAT), results in CMT1 with an early-onset demyelinating neuropathy with foot deformity and gait impairment. The reduced motor NCVs in patients with the I50del mutation indicate neuropathy in lower and upper limbs, but no defects in sensory nerves were observed (Geroldi et al., 2020).

In the present study, we describe the high-resolution crystal structures of the three recently discovered CMT-associated P2 variants; I50del, M114T, and V115A. The stability and lipid binding of these variants were investigated using circular dichroism (CD) spectroscopy, lipid vesicle aggregation assays, time-lapse imaging of supported lipid bilayers, and fatty acid binding assays. In addition, a room-temperature crystal structure of human P2 is reported through a joint neutron/X-ray refinement and used together with MD simulations to get further information on P2 flexibility.

## Results

P2 is a small β barrel protein of the FABP family (Fig 1). Previously, we studied the structure-function relationships in three CMT-linked variants: I43N, T51P, and I52T (Ruskamo et al., 2017). Now, three more disease variants have been reported: I50del, M114T, and V115A (Geroldi et al., 2020; Palaima et al., 2019). On the wild-type P2 (P2-wt) 3D structure (Majava et al., 2010), all six CMT-linked mutations are clustered close to one another (Fig 1).

**Fig 1.**
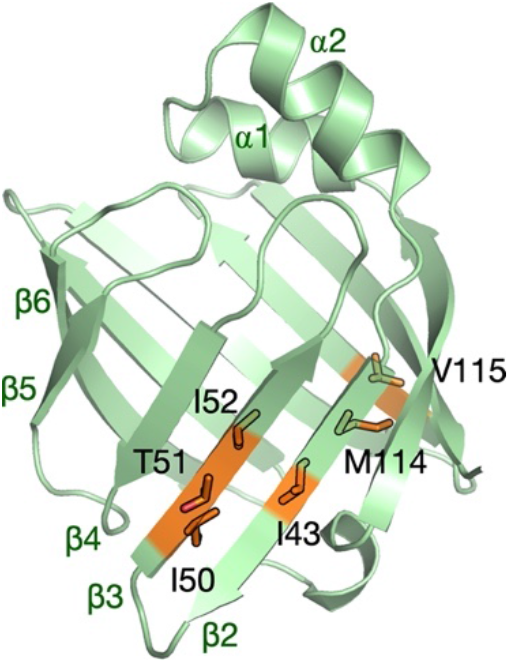
Overall structure of human P2. The locations of all known CMT-linked mutations on the protein structure have been indicated (orange) and labelled. Key secondary structure elements are indicated in green.

To elucidate the structure and function of the three recently discovered P2 variants, we expressed and purified V115A, M114T, and I50del. All the variants behaved well, and none of them showed signs of aggregation during purification or further analysis. We used both experimental and *in silico* techniques to study the effects of the P2 patient mutations on structure and function at the molecular level.

### The crystal structures of CMT-linked P2 variants

P2-I50del, M114T, and V115A were crystallised, and structures of all variants were refined at high resolution (Table 1). The asymmetric unit of I50del and M114T contained a single protein molecule, whereas that of V115A had two P2 molecules. The overall fold of all variants, with a β barrel covered by an α-helical lid, remained unchanged in comparison to P2-wt. In the V115A crystal structure, the α-helical lid and hinge regions of two molecules face each other. All structures contained a palmitate molecule bound inside the β barrel. The CMT mutation sites are situated at the bottom of the β barrel (Fig 1). Ile50 is located on strand β3, whereas M114T and V115A are located on strand β9 on the opposite side of the β barrel (Fig 2A).

**Table 1.** Data processing and structure refinement statistics. The processing statistics of the neutron dataset have been reported before (Laulumaa et al., 2015a).

|  |  |  |  |  |  |
| --- | --- | --- | --- | --- | --- |
| variant | I50del | M114T | V115A | P2-wt<br>X-ray | P2-wt<br>neutron |
| PDB ID | 7NSR | 7NRW | 7NTP | 7O60 |  |
| Data collection |  |  |  |  |  |
| Beamline | P11/PETRA III | P13/PETRA III | P11/PETRA III | rotating anode | LADI-III/ILL |
| X-ray wavelength (Å) | 1.033 | 0.976 | 1.033 | 1.54 | 3.0-3.9 |
| Space group | P 4<sub>1</sub> 2<sub>1</sub> 2 | P 4<sub>1</sub> 2<sub>1</sub> 2 | I 2 2 2 | P 4<sub>1</sub> 2<sub>1</sub> 2 | P 4<sub>1</sub> 2<sub>1</sub> 2 |
| Unit cell dimensions<br>a, b, c (Å) | 66.14, 66.14,<br>101.19 | 64.79 64.79<br>100.93 | 85.47 91.22<br>107.05 | 58.72 58.72<br>101.89 | 57.95<br>57.95<br>100.79 |
| α, β, γ (°) | 90, 90, 90 | 90, 90, 90 | 90, 90, 90 | 90, 90, 90 | 90, 90, 90 |
| Resolution range (Å) | 50-1.50 (1.54-1.50) | 50-2.00 (2.05-2.00) | 50-2.10 (2.15-2.10) | 15-2.00 (2.06-2.00) | 40-2.40<br>(2.53-2.40) |
| No. unique<br>reflections | 36645 (2657) | 15135 (1092) | 24729 (1814) | 12464 (891) | 4232 (363) |
| Completeness (%) | 99.9 (99.3) | 99.9 (99.7) | 99.7 (99.9) | 98.9 (96.1) | 60.1 (36.9) |
| Redundancy | 13.9 (13.7) | 13.8 (13.5) | 4.9 (5.0) | 9.7 (3.2) | 4.2 (2.2) |
| R<sub>sym</sub> (%) | 6.6/573.4 | 24.5 (326.7) | 7.0/254.6 | 6.0 (43.1) | 18.9 (30.4) |
| R<sub>meas</sub> (%) | 6.9 (595.8) | 25.5 (339.6) | 7.8 (285.0) | 6.3 (51.3) | - |
| <I/σI> | 17.6 (1.1) | 11.6 (1.2) | 11.2 (0.9) | 25.2 (3.0) | 6.1 (2.8) |
| CC<sub>1/2</sub> (%) | 99.9 (74.3) | 99.4 (70.3) | 99.8 (26.1) | 99.9 (84.9) | - |
| Wilson B (Å<sup>2</sup>) | 25.9 | 44.6 | 58.98 | 34.3 | - |
| Structure refinement |  |  |  |  |  |
| R<sub>cryst</sub>/R<sub>free</sub> (%) | 18.2/19.9 | 20.7/25.0 | 23.7/28.9 | 15.5/20.3 | 29.5/34.1 |
| RMSD bond lengths<br>(Å) | 0.022 | 0.012 | 0.0059 | 0.029 |  |
| RMSD bond angles<br>(°) | 1.880 | 1.1 | 0.808 | 2.1 |  |
| Molprobability score<br>(percentile) | 1.72 (67th) | 0.66 (100th) | 1.49 (98th) | 2.03 (71st) |  |
| Ramachandran<br>favoured/outliers<br>(%) | 97.7/0 | 100/0 | 96.2/0 | 97.7/0.8 |  |

**Fig 2.**
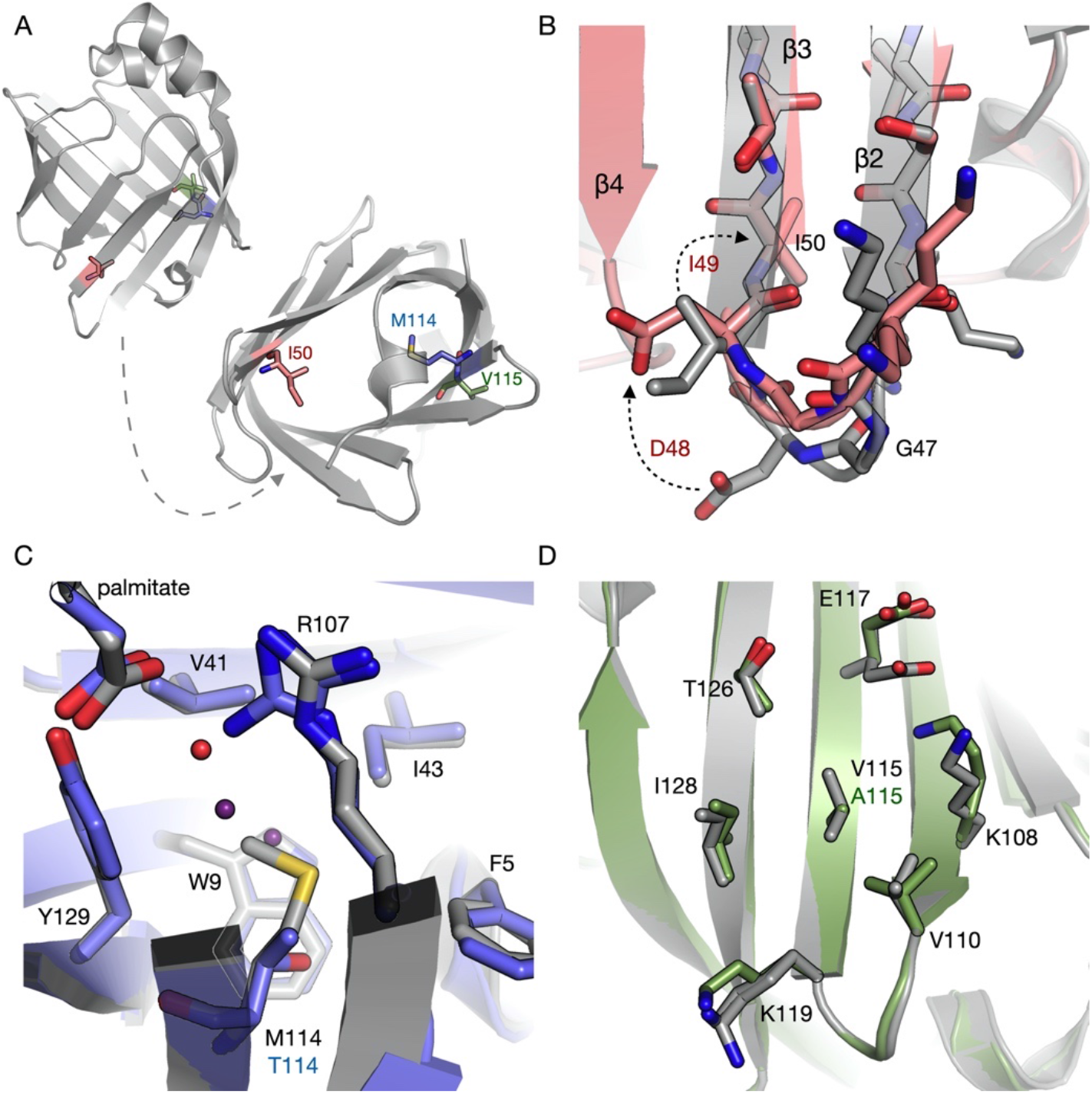
Crystal structures of new CMT variants of human P2. A. Overall view from the side and bottom of P2, highlighting the locations of Ile50 (pink), Met114 (blue), and Val115 (green). B. Comparison of P2-wt (grey) and I50del (pink). The deletion of Ile50 leads to Ile49 taking its buried position and a change of register in the β2-β3 loop. C. Comparison of P2-wt (grey) and M114T (blue). Arg107 is in a double conformation in M114T, and the cavity resulting from the mutation has two new water molecules (blue spheres). P2-wt has one structural water molecule nearby (red sphere). D. Comparison of P2-wt (grey) and V115A (green). Val115 is in a central position on the β sheet surface, and the mutation causes some rearrangements of nearby residues through altered van der Waals interactions.

The I50del mutation is located in the beginning of the β3 strand in the close proximity of the β2-β3 loop. The deletion shortens the loop and locally changes the sequence register, but it has no effect on the hydrogen bonding of the β sheet (Fig 2B). In both P2-wt and the mutant, the side chain of Asp48 interacts with the Lys66 side chain, located in the loop β4-β5. In P2-wt, Asp48 is situated near to the tip of the β2-β3 loop, while in I50del, this residue lies closer to the central region of the β barrel, in the beginning of strand β3. In P2-wt, Ile49 occupies this position.

In our crystal structures, both M114T and V115A have minor effects on the conformation or the hydrogen bonding of the surrounding amino acids (Fig 2C,D). Met114 points towards the fatty acid inside the β barrel, but does not directly interact with the bound ligand (Fig 2C). Arg107, located close to Met114 on the adjacent β strand (β10), interacts directly with the bound palmitic acid. This residue adopts two side chain conformations in the M114T structure, indicating additional space and flexibility in the mutant protein. In one of these conformations, the Arg107 guanidino group distance to the carboxyl group of palmitic acid is longer, possibly altering the fatty acid binding affinity of the M114T variant. Additionally, two extra water molecules are present in a cavity caused by the M114T mutation (Fig 2C). These water molecules form hydrogen bonds to Arg107 and Thr114 side chains, respectively.

Val115, on the other hand, is exposed and points outwards from the β barrel (Fig 2D); its side chain forms van der Waals interactions with neighboring residues. On the protein surface, Val115 is situated in the middle of a small strip of hydrophobic residues and charged residues. The V115A mutation slightly changes the conformation of these residues and thereby may somewhat affect the surface electrostatics of P2. Another unique feature in the V115A structure is observed in loop β3-β4 of chain A; the side chain of Phe58 has flipped outwards and the palmitic acid is shifted closer to the loop.

### Solution structures of P2 variants

As seen above, the crystal structures of all CMT-associated P2 mutants closely resemble P2-wt. However, in a study on earlier P2 mutations, we observed changes in the solution behaviour in two CMT variants, whereby they opened up in solution (Ruskamo et al., 2017). This opening may be a functional property of P2 during lipid ligand entry and egress, as well as lipid bilayer binding (Laulumaa et al., 2018; Ruskamo et al., 2020). Hence, we used synchrotron small-angle X-ray scattering (SAXS) to study, if the P2 variant solution structures differ from P2-wt.

All variants behaved well in the SAXS measurements, and the Guinier regions were linear, in line with a monodisperse monomeric sample. The SAXS scattering curves and distance distributions of P2-wt and disease variants were similar (Fig 3A,B); the radius of gyration (R_g_) and maximum distance (D_max_) remained unchanged (Table 2). The results confirm that, in line with the crystal structures, the solution structures of the CMT-linked variants are similar to that of P2-wt.

**Fig 3.**
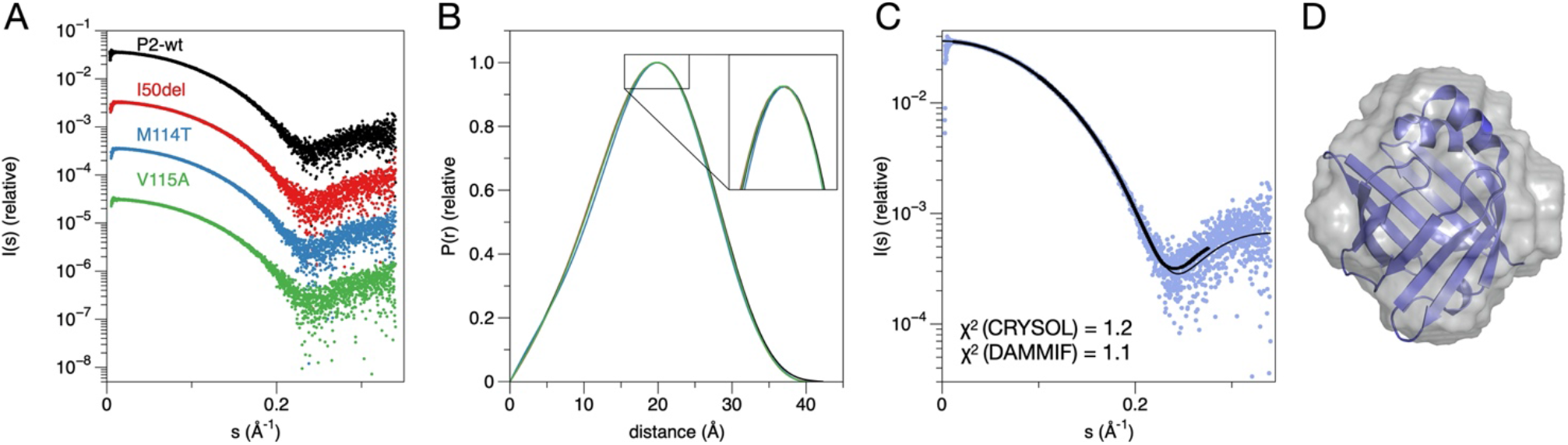
Solution studies on P2 mutants using SAXS. A. Scattering curves from a SEC-SAXS experiment. B. The distance distribution function indicates essentially identical structures in solution for all 4 samples. C. Fits of the crystal structure and a dummy atom model to the raw data. D. P2-wt crystal structure (blue cartoon) superimposed on the *ab initio* dummy atom model based on SAXS.

**Table 2.** SAXS parameters.

| sample | $R_g$ (Å) | $D_{max}$ (Å) | $V_{porod}$ (Å<sup>3</sup>) |
| --- | --- | --- | --- |
| P2-wtwt | 14.72 | 42.3 | 17400.90 |
| I50del | 14.65 | 40.3 | 17080.10 |
| M114T | 14.65 | 40.4 | 16829.60 |
| V115A | 14.63 | 40.2 | 15843.20 |

### Stability and folding

CD spectroscopy was used to compare the secondary structure content and thermal stability of the P2 variants (Fig 4). The shape of the CD spectra of the variants and P2-wt was similar (Fig 4A), but the intensity of the positive and negative peak maxima slightly varied; this may indicate slightly different average degrees of folding in solution or - more likely - minor errors in concentration.

**Fig 4.**
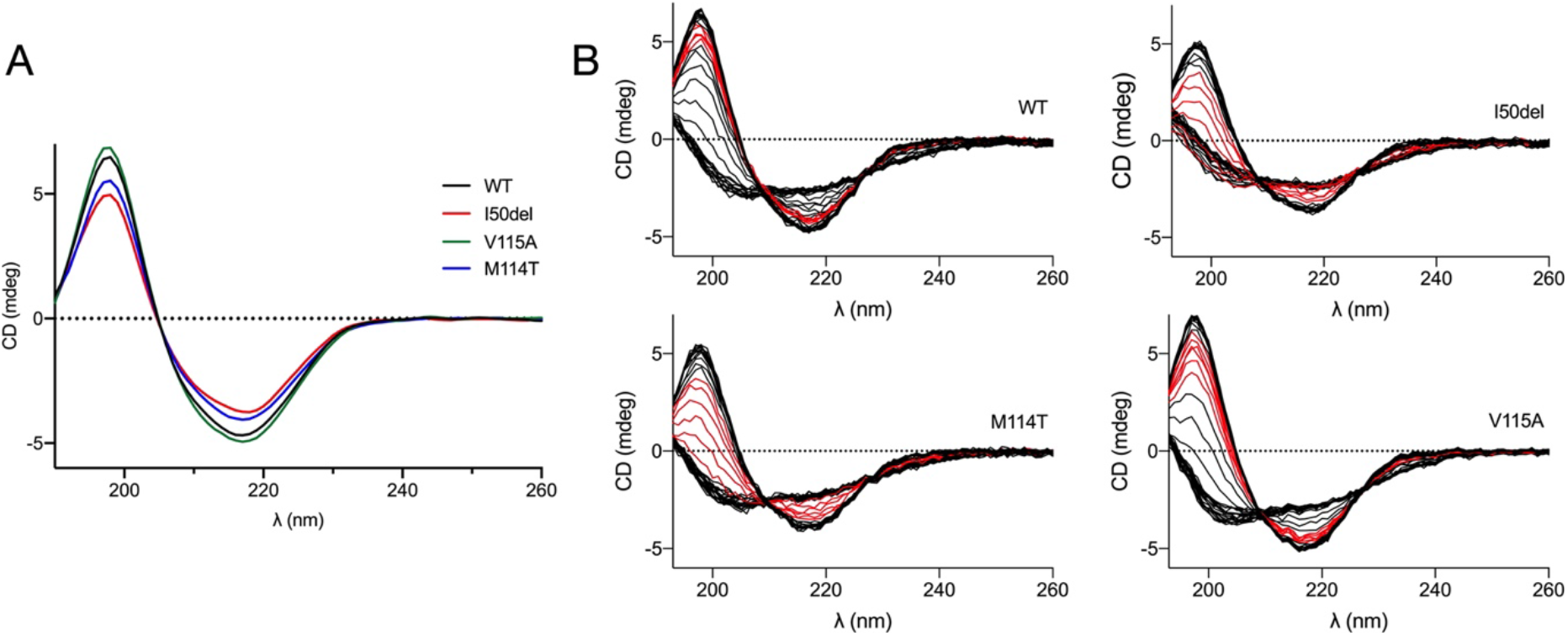
Folding and stability. A. CD spectra of P2-wt and the 3 variants. B. Assay of temperature stability. The measured CD curve is shown every 2 degrees upon heating, with the temperature range from +46 to +56 °C shown in red for each variant.

We studied thermal stability and measured the melting temperatures (T_m_) using CD. The T_m_ for P2-wt was +61.6 °C, whereas I50del showed much lower T_m_ (+46.3 °C) (Fig 4B, Table 3). The T_m_ of the missense variants M114T and V115A were +49.2 °C and +56.3 °C, respectively, also showing a decrease in thermal stability (Fig 4B). Thus, although the crystal and solution structures of the P2 disease variants were nearly unchanged compared to P2-wt, the stability of the CMT-associated variants is remarkably reduced and may affect the function of variants *in vivo*.

**Table 3.** Summary of structure and function of currently known human P2 disease variants. ΔT_m_ is given relative to P2-wt in each individual study.

| mutation | structural features | location | $\Delta T_m$ (°C) | fatty acid binding | membrane stacking | phenotype | references |
| --- | --- | --- | --- | --- | --- | --- | --- |
| I43N | open in solution | $\beta 2$, inwards | -17 | increased | unstable | Reduced NCVs (11-21 m/s), mild or moderate muscle weakness and atrophy, foot malformation, abnormal myelin, onion bulbs, neuronal abnormalities in zebra fish, reduced performance in rotarod test and reduced amount of large myelinated axons in transgenic mice | (Gonzaga-Jauregui et al., 2015; Hong et al., 2016; Ruskamo et al., 2017) |
| I50del | $\beta 2$-$\beta 3$ loop shorter | start of $\beta 3$, inwards | -16 | normal | normal | Reduced NCVs (~20-30 m/s), equinus foot deformity and gait impairment | (Geroldi et al., 2020), this study |
| T51P | open in solution; tends to aggregate | $\beta 3$, outwards | -24 | increased+ | decreased | Reduced NCVs (~11 m/s), severe muscle weakness and atrophy, foot malformation | (Motley et al., 2016; Ruskamo et al., 2017) |
| I52T | loss of H bonds | $\beta 3$, inwards | -13 | increased | unstable | Reduced NCVs (14-21 m/s), muscle weakness and atrophy in lower and upper limbs, reduced density of myelinated axons and thickness of myelin, onion bulbs | (Motley et al., 2016; Ruskamo et al., 2017) |
| M114T | extra water molecules, increased dynamics | start of $\beta 9$, inwards | -13 | increased | normal | Reduced NCVs (<20 m/s), frequent falls, muscle weakness, severe demyelination and secondary axonal degeneration | (Palaima et al., 2019), this study |
| V115A | altered surface properties | $\beta 9$, outwards | -6 | increased | normal | Mostly normal NCVs, clumsiness, foot deformity, variable and very mild demyelination and focal pattern of distribution along peripheral nerves | (Palaima et al., 2019), this study |

### Molecular dynamics simulations

The dynamics of wild-type and mutant P2 were studied with >1-μs MD simulations in water. The starting points for the simulations were the individual crystal structures. The RMSF plots were analysed to detect local differences in dynamics coupled to the mutations. Furthermore, plots of R_g_ and the opening of the P2 barrel were followed through the simulation (Fig 5).

**Fig 5.**
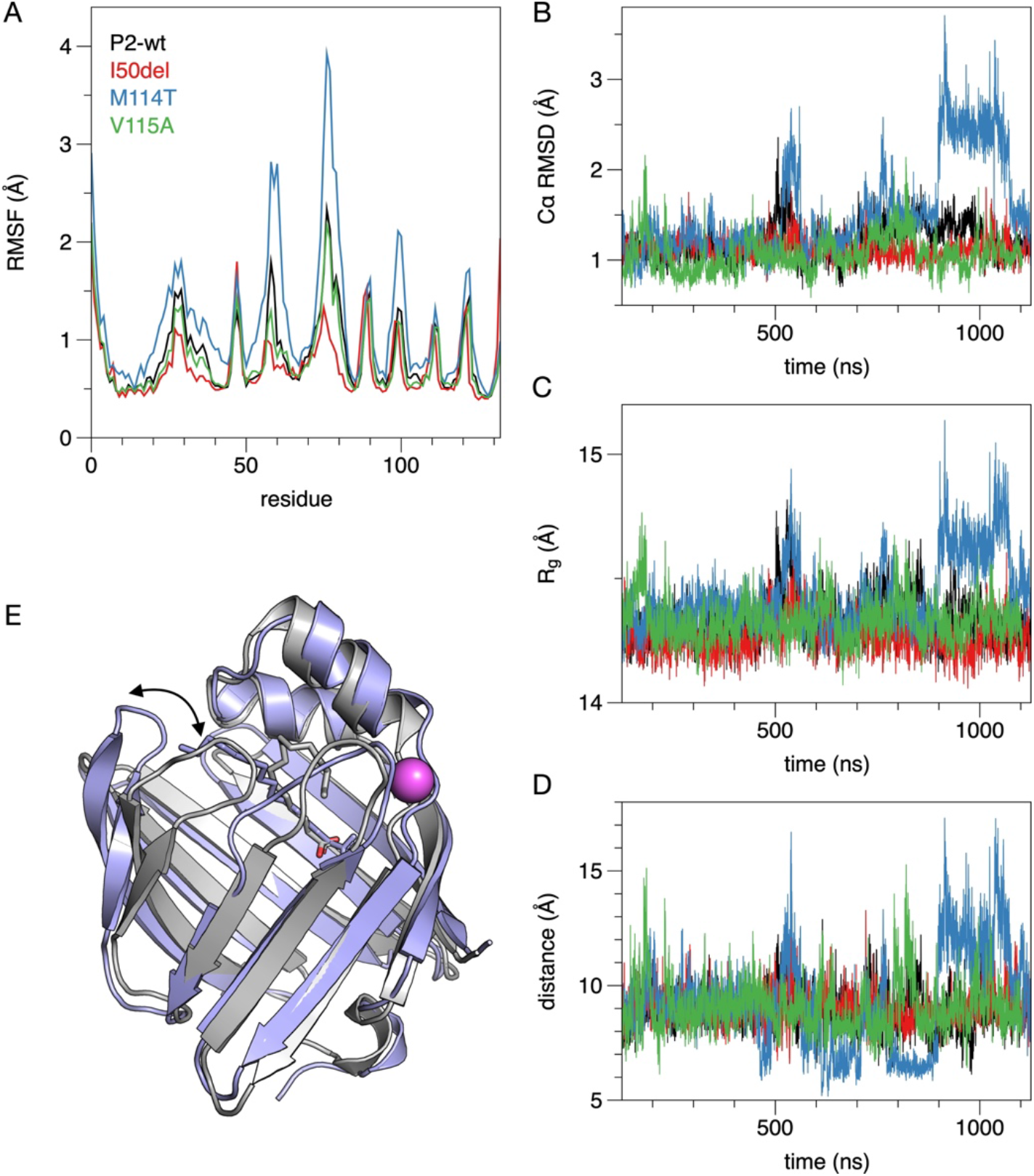
MD simulation of P2-wt and CMT variants. A. RMSF plots during the simulation. B. Cα RMSD compared to the starting structure. C. R_g_ during the simulation. D. Opening of the β barrel between strands β4 and β5, indicated by the distance between the Cα atoms of residues 61 and 74 and linked to the movement of the β5-β6 unit away from the helical lid. E. Snapshot of the opened M114T structure at a time point of 920 ns (blue), overlaid with the crystal structure (grey). The anion binding site is indicated with a magenta sphere, and the bound palmitate molecules are shown as sticks. Opening of the β5-β6 unit is indicated by the arrow.

The results indicate, as expected, highest mobility for the loop regions of the β barrel, especially in the β5-β6 region around residues 75-80. The M114T mutant had higher RMSF values, indicating an overall effect on protein dynamics by this non-conservative mutation in a buried residue side chain. I50del, on the other hand, was less dynamic than the other variants, which correlates with its short β2-β3 loop.

### Crystal structure of perdeuterated human P2 at room temperature

All published human P2 crystal structures thus far have been determined from cryo-cooled crystals at 100 K. In order to gain additional insight into P2 structure and dynamics, its crystal structure was here determined at room temperature. To this end, crystals of perdeuterated human P2 (Laulumaa et al., 2015a) were subjected to both neutron and X-ray diffraction data collection at room temperature. A joint refinement using both datasets was then carried out, and the room temperature crystal structure was compared to the structure at 100 K as well as to the above MD data.

B factor analysis reveals that while the overall shape of the B factor plot is similar, the β5-β6 hairpin loop is more dynamic at room temperature (Fig 6A,B). This loop corresponds to the strands in the β barrel, which present a large conformational change upon β barrel opening (Laulumaa et al., 2018; Ruskamo et al., 2020, 2017). When P2 binds to a lipid bilayer, the β5-β6 unit flaps open and interacts directly with the membrane surface lipid headgroups (Ruskamo et al., 2020). The RT crystal structure is in line with high flexibility of the portal region and especially strands β5-β6. Examples of electron and nuclear density maps of the RT structure of P2 are shown in Fig 6C-E.

**Fig 6.**
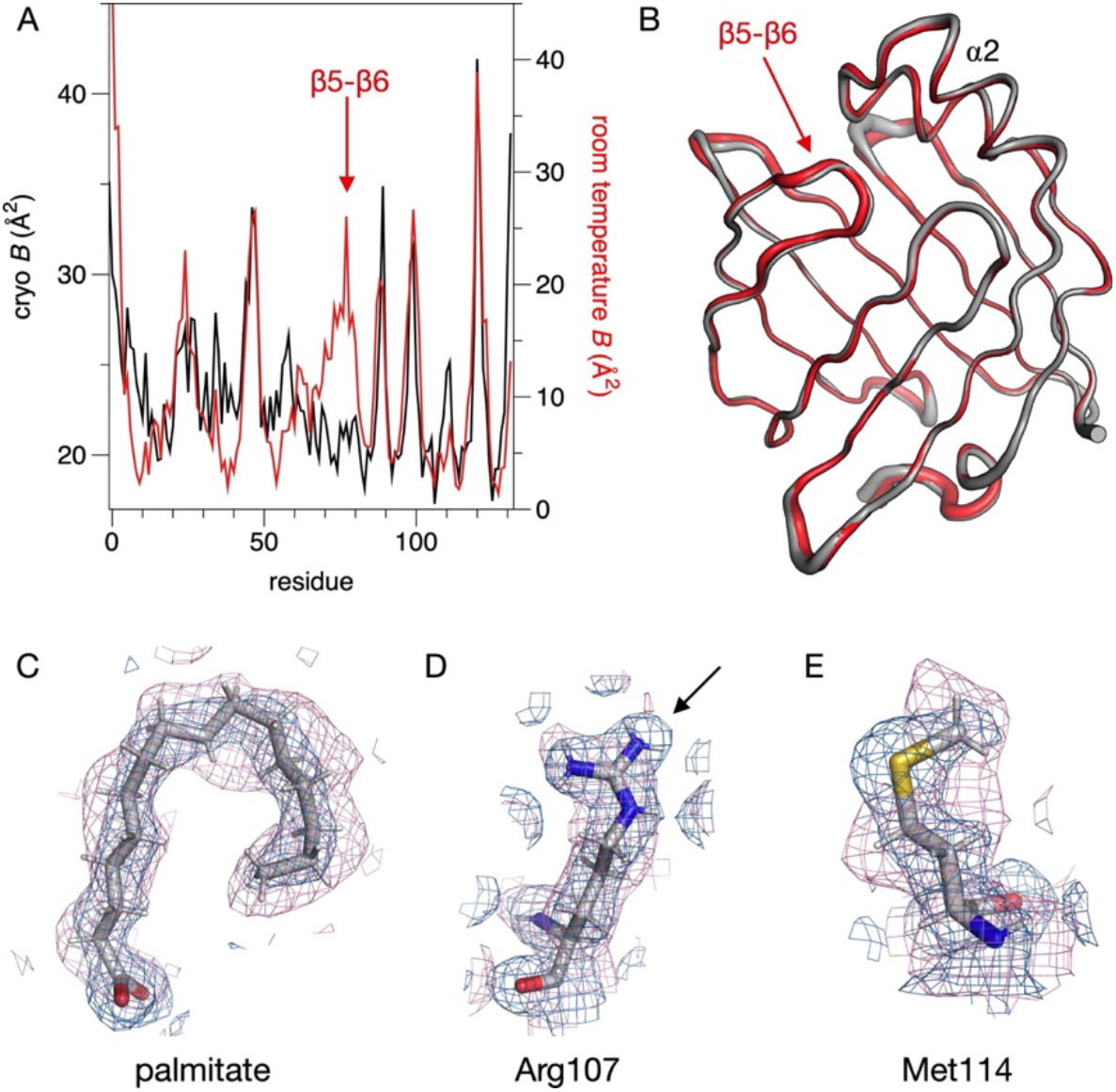
B factor-based analysis of dynamics in the crystal state. A. Average B factor plot of human P2 at 100 K (black) and at room temperature (red). B. Overlay of the cryo (gray) and RT (red) structures, with the thickness of the ribbon reflecting the B factor. C-E. Electron (blue) and nuclear (pink) density maps of selected regions of the structure. The 2F_o_-F_c_ maps are both contoured at 1 σ. The arrow in D indicates lack of nuclear density for part of the Arg107 guanidinium group. Arg107 was previously shown (Laulumaa and Kursula, 2019; Ruskamo et al., 2014) to be deprotonated in cryocooled crystals using ultrahigh-resolution X-ray crystallography.

Comparing to the MD simulation data, the RT structure complements the story. In MD simulations, especially for M114T (Fig 5D,E), the β5-β6 flap is the most mobile segment, while its B factors are low in cryo-cooled crystals. A clear increase in the mobility of the β5-β6 segment is seen in the RT crystal structure, pointing towards a functionally relevant difference between RT and cryo-cooled crystal structures.

### Bioinformatics analyses for the mutations

The sequence conservation of P2 among selected vertebrates was studied with multiple sequence alignments (Fig 7A), and the conserved residues were mapped onto the P2 crystal structure (Fig 7B,C). The conservation has interesting patterns; essentially every second residue on the β strands is conserved, corresponding to inward-pointing side chains. In addition, the inside of helix α1 and the outside of helix α2 are conserved. The latter is hydrophobic and expected to be embedded in the bilayer core upon membrane binding (Ruskamo et al., 2020, 2014), while helix α1 interacts with the bound fatty acid. Furthermore, the loop β2-β3 and strands β4-β5 are conserved, as is Pro39, which is important for P2 dynamics and membrane interactions (Laulumaa et al., 2015b; Ruskamo et al., 2014).

**Fig 7.**
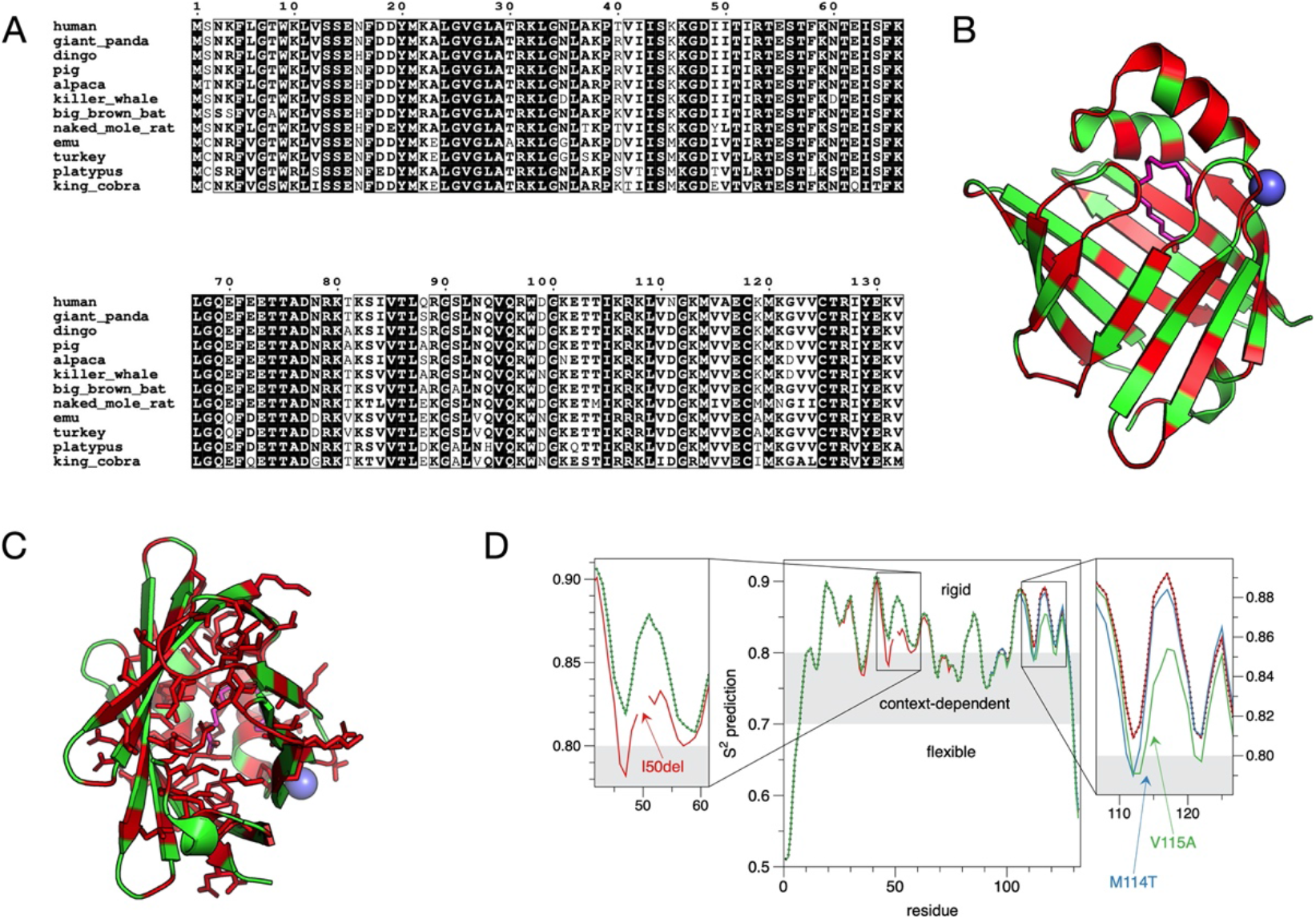
Conservation mapped onto structure. A. Sequence alignment of P2 from 12 selected vertebrates. Fully conserved residues are shaded on black background. B. Mapping of the fully conserved residues (red) onto the P2 structure. The sphere indicates the anion-binding site and the hinge region, and the palmitate inside the β barrel is shown as sticks. C. View from the β barrel bottom shows how the side chains of conserved residues in the barrel mainly point inwards. D. Sequence-based dynamics prediction suggests that all CMT mutations cause increase in local flexibility.

A sequence-based prediction of rigidity was carried out using DynaMine. All three mutations were predicted to cause an increase in local flexibility of the P2 structure (Fig 7D). The result is similar to that observed with the other CMT-linked variants of P2 (Laulumaa et al., 2018), which suggests similar molecular mechanisms for the different disease variants.

### Molecular functions of disease variants

One of the suggested main functions of P2 in PNS myelin is to glue stacked lipid bilayers together. To investigate the membrane stacking ability of P2 variants, they were subjected to a lipid vesicle aggregation assay (Fig 8A), in which the vesicle aggregation induced by P2 is monitored as a change in a solution turbidity. Negatively charged model membrane systems of 1,2-dimyristoyl-*sn*-glycero-3-phosphocholine (DMPC) : 1,2-dimyristoyl-*sn*-glycero-3-phosphorylglycerol (DMPG) vesicles were used. No significant differences were observed in turbidity between P2-wt and mutants. All variants gave the strongest signal at 10 μM, which corresponds to a molar P/L ratio of 1:50. While one can expect highest stacking activity when each membrane surface is 50% or less saturated, it is likely that higher protein concentrations will saturate individual membranes and prevent stacking.

**Fig 8.**
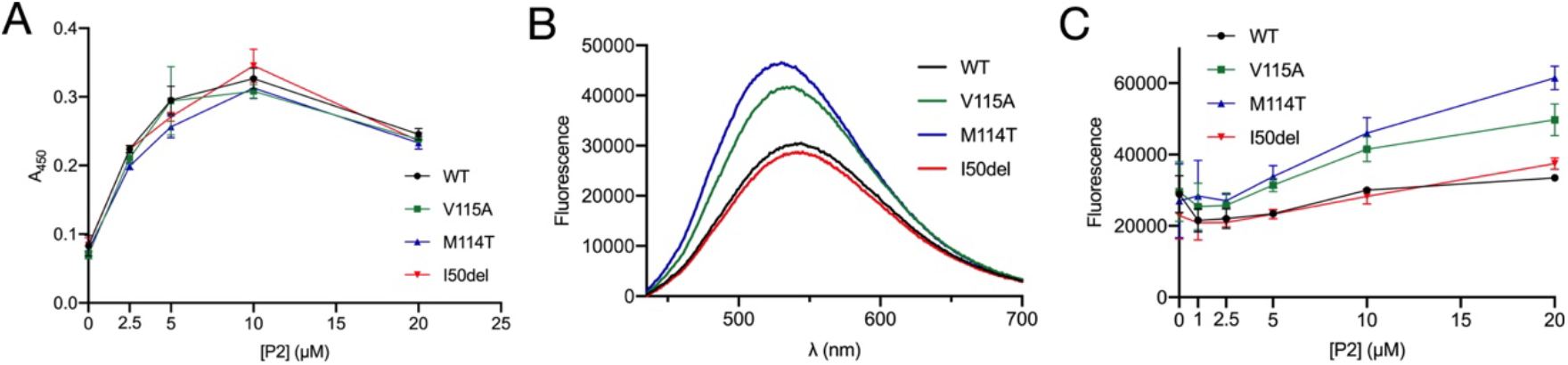
Binding to membranes and fatty acid ligands. A. Turbidimetric analysis of DMPC:DMPG lipid vesicle aggregation induced by P2 variants. The figure shows the average of three measurements. B. Fluorescence spectra of DAUDA in the presence of 10 μM P2. C. Titration of DAUDA with P2, fluorescence measured at 535 nm. All points were measured in triplicate.

11-dansylaminoundecanoid acid (DAUDA) is an environment-sensitive fluorescent fatty acid derivative probe, whose fluorescence emission spectrum changes upon interaction with a protein. No clear difference was seen in the spectrum of I50del compared to P2-wt (Fig 8B,C). In contrast to I50del, both M114T and V115A showed a change in the fluorescence peak intensity at 535 nm compared to P2-wt (Fig 8C). The fluorescence intensity of DAUDA with M114T and V115A increased, indicating enhanced binding of the fatty acid probe to these variants. This can be an indication of increased protein dynamics coupled to less saturation with bound *E. coli* fatty acids in the mutant protein preparation, being reminiscent of previously studied P2 variants (Laulumaa et al., 2018, 2015b; Ruskamo et al., 2017, 2014).

### Myelin protein P2 induces membrane blebbing

Time-lapse fluorescence microscopy was employed to explore the effects of human P2 (P2-wt, P2-M114T, P2-I50del, and P2-V115A) on planar double supported membrane patches. An illustration of the structure of a double supported membrane patch is shown on the top of Fig 9. The isolated bilayer patches with free edges typically have diameters in the range of 50-100 μm. The model membrane composition was 90% 1,2-dioleoyl-*sn*-glycero-3-phosphocholine (DOPC), 10% 1,2-dioleoyl-*sn*-glycero-3-phospho-L-serine (DOPS), and the membranes were hydrated in 10 mM Tris buffer with 2 mM Ca^2+^. Myelin P2 proteins were added to the fluid cell from a concentrated solution to a final bulk concentration of 33.3 μM in the cell. After addition, the proteins reach the membrane patch on a timescale of 30-60 s.

**Fig 9.**
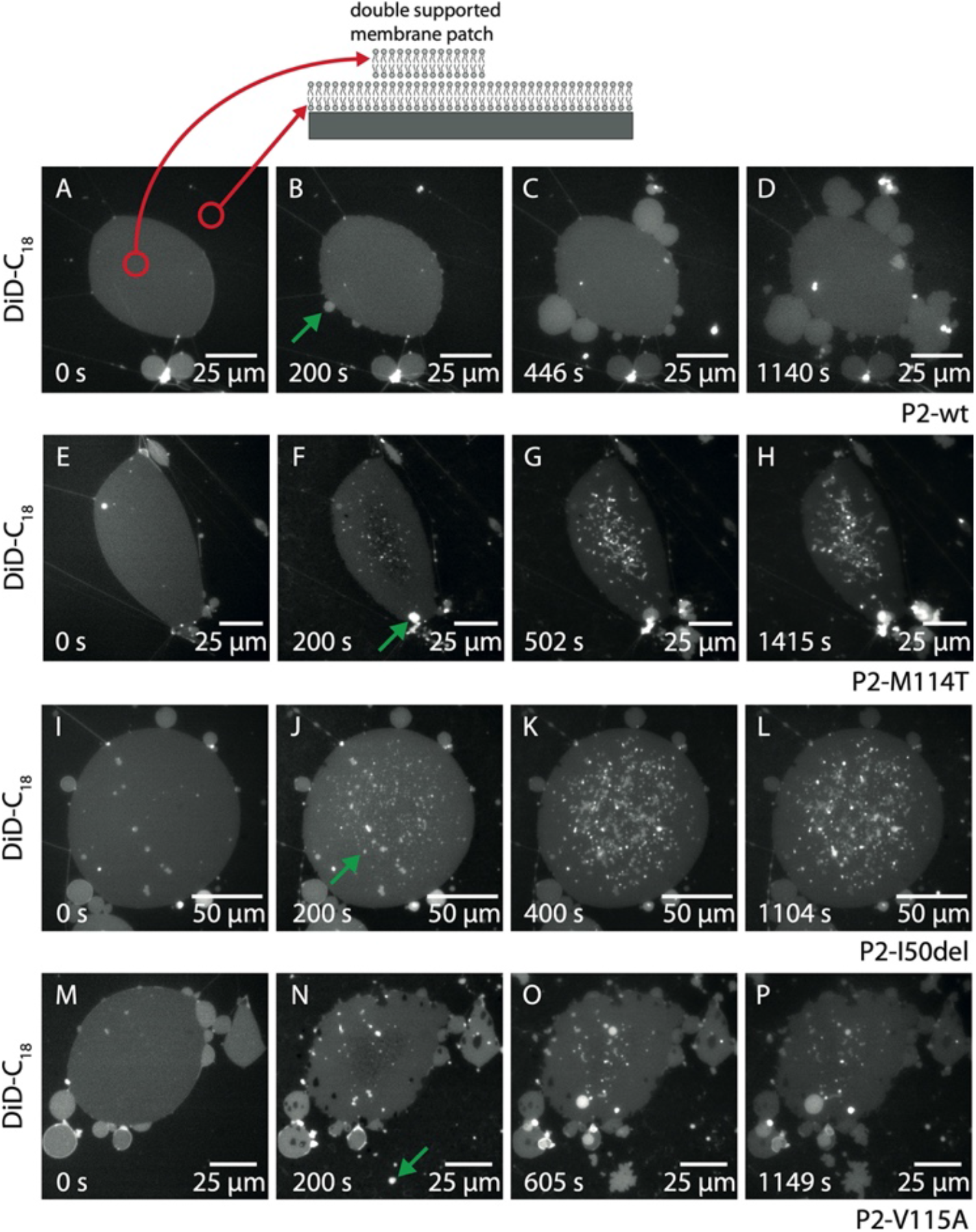
Time-lapse fluorescence sequences showing membrane blebbing induced by myelin protein P2. Schematic of a double supported model membrane patch (**top**). P2wt induces considerable blebs mainly initiated at the edges of the double supported membrane patch (**A-D**). Upon exposure to P2 mutants P2-M114T (**E-H**), P2-I50del (**I-L**) and P2-V115A (**M-P**) a significant amount of primarily smaller blebs emerges from the membrane patch. The green arrows mark examples of membrane blebs. All proteins were at 33.3 μM concentration. N=4 repetitions.

After exposure to P2-wt, round vesicular structures were observed to emerge from the membrane surface as viewed in Fig 9B-D. We refer to these vesicular structures as membrane blebs, since they resemble structures from previous studies (Boye et al., 2018). The membrane blebs emerge after ~2 min, primarily at the edges of the double supported membrane patch, and continue to grow until ~20 min.

For comparison, we studied the effects induced by three P2 mutants; the results are shown in Fig 9E-H (P2-M114T), Fig 9I-L (P2-I50del), and Fig 9M-P (P2-V115A). Time-lapse videos of the experiment are additionally available as Supplementary Information. The P2 mutants primarily induced a large number of smaller membrane blebs, mainly emerging from within the double supported membrane patch, and not at the edges as observed for P2-wt. The small blebs are initiated on a similar timescale as observed for the larger blebs induced by P2-wt. That is, the curvature effects induced by the P2 mutants are smaller, but they occur on a similar timescale as P2-wt.

## Discussion

P2 is a major component of PNS compact myelin (Trapp et al., 1984, 1979) and plays a role in lipid homeostasis of Schwann cells and in peripheral myelin remodeling (Stettner et al., 2018; Zenker et al., 2014). P2 mutations are inherited in an autosomal dominant manner and lead to demyelinating CMT1 with varying severity (Gonzaga-Jauregui et al., 2015; Hong et al., 2016; Motley et al., 2016; Punetha et al., 2018). Recently, three novel CMT1-associated P2 mutations (I50del, M114T, and V115A) were discovered (Geroldi et al., 2020; Palaima et al., 2019), but no experimental data on the structure or biophysical and biochemical characteristics of the corresponding protein products have been available. Our work provides new details of P2 function and may help to elucidate its function in myelin maintenance and remyelination as well as shed light on molecular mechanisms of CMT.

### Disease variants show reduced stability despite similar structure

P2 has a compact and stable β barrel structure similar to other members of the FABP family. All residues currently known to be affected by disease mutations are conserved among most mammalian species. Nevertheless, none of the new mutations had major effects on the crystal structure of P2. Similar results were obtained earlier with P2 I43N, T51P, and I52T disease variants (Ruskamo et al., 2017).

Since crystallisation may favour the most stable conformation and hide functionally relevant differences, the structures of P2 variants were studied in solution. SAXS analysis revealed no differences in conformation, indicating correct folding and a closed β barrel of the variants in solution. Hence, all variants studied here closely resemble the crystal structure in solution. Previously, we observed an altered X-ray scattering pattern and a more open conformation of the T51P disease variant in solution, which was linked to its reduced stability and altered ligand-binding properties (Ruskamo et al., 2017). Such differing solution dynamics may further be linked to both fatty acid ligand binding and membrane bilayer stacking.

The most drastic consequence of the P2 disease mutations studied here was observed in the thermal stability of the variants. Similarly, all previously studied CMT1-associated mutations showed 13 to 24 degrees reduced thermal stability compared to P2-wt, T51P being the most unstable (Ruskamo et al., 2017). I50del and M114T had 15 and 12 degrees lower thermal stability indicating the T_m_ change in the same range with the earlier studied I43N and I52T variants (Ruskamo et al., 2017). V115A had a smaller effect on protein stability; however, a drop in T_m_ of several degrees correspond to a large energetic difference, which is surprising considering the kind and location of this mutation on the outside of the barrel. Interestingly, the family carrying the V115A mutation did not show NCVs typical for CMT1 patients and suffered milder symptoms compared to patients with other P2 mutations (Palaima et al., 2019). This link between P2 stability and clinical features of the CMT1 patients is an important topic for future research.

### Fatty acid and membrane bilayer binding

P2 binds to lipid membranes using two opposite faces of the protein, leading to membrane stacking and formation of multilayered myelin-like assemblies (Knoll et al., 2014; Ruskamo et al., 2020, 2014; Suresh et al., 2010). Additionally, P2 binds and transports fatty acids into and from lipid membranes (Zenker et al., 2014). Here, we observed an enhanced binding of both point mutant variants to DAUDA, whereas I50del had no effect. These results resemble the ones obtained earlier with I43N and I52T mutants (Ruskamo et al., 2017) and may indicate changes in fatty acid binding dynamics, which could influence lipid metabolism of Schwann cells and myelin membrane lipid composition. On the other hand, we did not observe any changes in lipid vesicle aggregation/membrane stacking properties of the disease variants. The results are compatible with a scenario, in which the recombinant P2 variants are correctly folded, but dynamics of the barrel opening and portal region are altered, which may be coupled to less fatty acid binding in the mutant protein preparations - leading to apparently higher fatty acid binding *in vitro*.

Myelin protein P2 binds to membranes and induces spontaneous stacking into multilayers (Knoll et al., 2014; Ruskamo et al., 2020, 2014). Furthermore, in these complexes, both the P2 protein and the lipids become less dynamic, indicating a synergistic stabilisation of proteolipid multilayers induced by the presence of P2 (Knoll et al., 2014, 2010; Ruskamo et al., 2020, 2014). In time-lapse experiments on membranes, all P2 variants induced blebbing. However, while the time scales were similar, all three CMT variants induced morphologically different blebs, whereby smaller membrane structures emerged upwards from the membrane plane. P2-wt induced larger blebs at the edges of the bilayer island. The blebbing effect found in this study is in line with previous results on membrane stacking, since blebbing can also be thought of as the formation of additional membrane layers. These functional aspects are central in beginning to understand the role membrane curvature plays in myelin-like membrane stacking, as common model systems use planar membranes or large vesicles.

In an earlier study, annexin A1 and annexin A2 both induced membrane blebbing on a timescale of <1 min (Boye et al., 2018). In comparison, the timescale of P2-induced blebbing is somewhat slower, as the effect is initiated ~2 min after the protein has reached the membrane patch. For P2-wt, the total area of the emerging blebs is qualitatively similar to the effect reported for the annexins (Boye et al., 2018), while the large number of smaller blebs induced by the P2 mutants has not previously been observed in such assays. The myelin membrane is highly curved, and since blebbing is a membrane curvature effect, the curving induced by myelin P2 proteins could be related to the curved myelin membrane. Given the somewhat different morphology of blebbed membranes, the mutated P2 variants may function differently from P2-wt between two curved membranes and in the presence of the other PNS major dense line proteins MBP and P0 - a system that is very challenging to study *in vitro*.

### Comparison to previously characterised P2 variants

Our work complements structure/function studies of currently known CMT-associated mutations in the *PMP2* gene. Table 3 presents an overview of the main points of current and earlier results on these variants at the protein level. While some of the mutations cause more severe impairment in the protein properties, they essentially all behave similarly. All P2 disease mutations cause a large decrease in protein thermal stability, and most of them affect dynamics and fatty acid binding. Not all have an effect on membrane stacking based on turbidimetric assays, and the work in this study shows that the mutations do not inhibit membrane blebbing caused by P2.

Some correlations can be observed between disease severity in patients and the biophysical properties of the individual mutant proteins (Table 3). The heat stability of V115A is affected much less than for the other mutants; patients having this variant have a very mild disease, with nearly normal NCVs (Palaima et al., 2019). On the other hand, M114T is one of the more structurally drastic mutations, having a non-conservative replacement of a buried side chain. The patients with M114T present severe demyelination (Palaima et al., 2019). T51P is the most unstable P2 mutant protein characterised, which can structurally be explained by the insertion of a Pro residue into a β strand (Ruskamo et al., 2017). Patients with T51P have severe drop in NCV, coupled to muscle weakness and atrophy (Motley et al., 2016). While these observations may hint towards more details of disease mechanisms, at the moment they must be treated as possible correlations. Similar apparent correlations between mutant protein properties and CMT disease phenotype were previously observed for the cytoplasmic tail of myelin protein P0 (Raasakka et al., 2019), which lies in the same compartment in the PNS myelin major dense line as P2. The fact that the thus far observed CMT mutations in P2 cluster to the same 3D region may suggest a mutation hotspot carrying residues important for P2 structure or function.

Other mutations have been introduced into P2 over the years for structural and functional studies. Three of the most interesting ones include L28D, P39G, and F58A (Laulumaa et al., 2018, 2015b; Ruskamo et al., 2020, 2014). Leu28 localizes at the tip of the α helical lid. In a cellular system, the L28D mutation inhibits the formation of cytoplasmic membrane domains observed with P2-wt, as well as revokes the increased melanoma cell invasion induced by P2-wt overexpression (Graf et al., 2019; Ruskamo et al., 2014). Additionally, *in silico* modeling predicts the embedding of Leu28 deep into the lipid bilayer during membrane interaction (Ruskamo et al., 2014). Pro39 is found in the hinge region of the lid and P39G is a generally more active P2 variant in all experiments, such as membrane stacking, fatty acid binding, and dynamic analyses (Laulumaa et al., 2018). Phe58 is located in the portal region of P2 and is involved in the control of the opening of the β barrel structure and membrane binding. This portal Phe residue is conserved in most FABPs and has a crucial role in FABP ligand binding and lipid transport (Laulumaa et al., 2018; Ruskamo et al., 2014; Simpson and Bernlohr, 1998). In case of the mutations affecting directly fatty acid-binding residues (R107E, R127E, and Y129F), neither mutated proteins nor effects on cell viability were detected (Graf et al., 2019).

### Insights into structural dynamics in P2

Conformational changes are likely to be central to the function of P2, both in lipid transport and in membrane stacking. The dynamics of the portal region are important in the entire FABP family, allowing for ligand entry and egress (Friedman et al., 2006; Ragona et al., 2014). The division of the FABPs into two groups based on lipid transport mechanisms (Storch, 1993; Thumser and Storch, 2000) gives further insight into the role of the portal region. P2 is a FABP with a collisional ligand transfer mechanism.

A common property of all disease mutant variants of human P2 is the decreased heat stability of the fold; this may then lead to abnormal fatty acid and lipid bilayer binding. This trend is continued with the mutations studied here. The identified disease variants of P2 localise to two small clusters in the 3D structure, on opposite sides of the β barrel structure (Fig 1). The location could be a sign of a hotspot for correct folding and/or functional dynamics related to membrane and lipid ligand binding. The mutations are far from the portal region, but P2 being a small protein, altering the dynamics and flexibility of one end of the β barrel could well have large effects on the opening of the portal region at the other end. Simulations close to lipid membrane surfaces would give additional information on functional dynamics of the P2 mutants. The fact that membrane stacking properties are not much affected by the mutations indicates the surface properties of the mutant variants *per se* are similar.

For an insight into human P2 dynamics, we combined MD simulations with room temperature crystallography. The crystal structure at room temperature, compared to the structure at 100 K, has high B factors in the β5-β6 loop, which is the most mobile part of P2, when the β barrel opens up. Simulations have reproducibly indicated this opening (Laulumaa et al., 2018, 2015b; Ruskamo et al., 2017), and we could detect it for the earlier studied P2 CMT variants in solution using synchrotron SAXS (Ruskamo et al., 2017). Hence, functionally relevant dynamics can be revealed by room temperature crystallography in comparison to cryocooled crystals.

### What is the function of P2 in humans?

Historically, P2 was characterised as an abundant component of the PNS myelin sheath; intriguingly, it is not present in all myelin sheaths, however (Trapp et al., 1979), and the amount of P2 varies between species (Greenfield et al., 1973; Singh et al., 1978). P2-deficient mice had a very mild phenotype, with effects on lipid homeostasis in the PNS at periods of active myelination (Zenker et al., 2014). The mutant mice also indicated a role for P2 in mouse PNS remyelination (Stettner et al., 2018). The identification of several *PMP2* point mutations in recent years in CMT families (Geroldi et al., 2020; Gonzaga-Jauregui et al., 2015; Hong et al., 2016; Motley et al., 2016; Palaima et al., 2019; Punetha et al., 2018) has shed more light on P2 as being important for the correct formation and maintenance of human PNS myelin. All these conclusions have been based on the assumption that P2 is more or less specific to myelinating PNS Schwann cells in vertebrates in general, and even more so in mammals.

Recently, several studies have highlighted strong expression of P2 in human astrocytes, while P2 is essentially missing in mouse astrocytes (Cai et al., 2015; Kelley et al., 2018). This observation has been linked to the size regulation of astrocytes, and expression of P2 in mouse astrocytes increased their size (Kelley et al., 2018). These results have implications for understanding the function of P2; mouse models will not be informative in relation to its function in human astrocytes.

Furthermore, as human PNS myelin is much thicker than that of mice, mouse models may not give a complete view of human P2 function even in PNS Schwann cells, given the difference in lifespan and the requirements this brings to proteins in stable biostructures, such as the myelin sheath. Myelin proteins are among the most long-lived proteins in vertebrates (Fornasiero et al., 2018; Toyama et al., 2013). These are important pointers towards critical analysis of mouse models in general - in this case for a single, apparently rather mundane but stable protein, which could have more widespread implications. In other words, P2 may be more important for humans, and possibly other large vertebrates, than small mammals, such as the mouse. This importance of P2 is not necessarily restricted to PNS myelin, and it may extend to CNS astrocytes.

### Conclusions

In current and earlier work, we have shown that the CMT-associated variants of human P2 have similar properties to each other. While crystal structures of all disease variants are nearly identical to P2-wt, with minor differences in local hydrogen bonding, the stability of the variants in solution and their function in lipid binding are affected. As P2 has been thought to function in both lipid transport and membrane multilayer stacking, different functional aspects could be affected by these properties. A functional, stable P2 could be more important in humans than in mouse models, and in addition to the importance of unraveling the role of P2 in human PNS myelin, the molecular function of P2 in human astrocytes, large cells with strong membrane synthesis, deserves further study.

## Materials and methods

### Mutagenesis

A construct containing human P2 with an N-terminal His-tag and Tobacco Etch virus (TEV) protease cleavage site cloned into the pTH27 vector (Majava et al., 2010) was used to express P2-wt and as a template plasmid in mutagenesis to produce the I50del, M114T, and V115A variants. Primers to generate P2 mutations were purchased from Eurofins Genomics (Supplementary Table S1). Mutagenesis was carried out using the QuikChange Site-Directed Mutagenesis protocol (Agilent) and Phusion^®^ High-Fidelity DNA Polymerase (New England Biolabs). Constructs were validated by DNA sequencing analysis.

### Protein expression

Proteins were expressed in *Escherichia coli* Rosetta (DE3) strain in ZYM-5052 autoinduction medium with 100 μg/ml of ampicillin at +18 °C for 44 h (Studier, 2005). The cells were harvested and suspended in a lysis buffer (300 mM NaCl, 10 mM imidazole, 50 mM HEPES pH 7.5, 1 mM DTT).

### Protein purification

For protein purification, cells were lysed by sonication and insoluble materials were pelleted by centrifugation (51000 x g, 30 min, at + 4 °C). The soluble fraction was mixed with the HisPur Ni-NTA Resin (Thermo Fisher Scientific) at + 4 °C for 2 h. The resin was washed two times with washing buffer (300 mM NaCl, 40 mM imidazole, 50 mM HEPES pH 7.5, 1 mM DTT) using centrifugation (300 x g, 4 min, at +4 °C). Then, the samples were transferred into a gravity-flow column and further washed with 100 ml of washing buffer. The samples were incubated for 5 min and eluted with elution buffer (300 mM NaCl, 300 mM imidazole, 50 mM HEPES pH 7.5, 1 mM DTT). To cleave the His tag, 40 μM of recombinant TEV protease was added. Imidazole was removed by dialysis through 6-8 kDa MWCO dialysis tubing (SpectraPor) against dialysis buffer (300 mM NaCl, 20 mM HEPES pH 7.5, and 1 mM DTT) at +4 °C for 17 h. TEV and the cleaved His tag were removed with a reverse immobilized metal affinity chromatography step using the HisPur Ni-NTA and the dialysis buffer. Proteins were further purified with a size-exclusion chromatography (SEC) using the dialysis buffer and a Superdex 75 pg 16/600 column (GE Healthcare). The proteins were concentrated with Amicon Ultra 15, MWCO 10 kDA protein concentrator (Merck Millipore) to the final concentration of 9-10 mg/ml.

### Crystallisation and structure determination

P2 mutants were crystallised at + 20 °C using the sitting drop vapour diffusion method. The concentrations of the I50del, M114T, and V115A mutants were 9.5, 9.5 and 10 mg/ml, respectively. M114T was crystallised in 2.4 M sodium malonate, pH 7.0. I50del was crystallised in 1.9 M sodium malonate, pH 6.4. V115A was crystallised in 2.2 M sodium malonate, pH 7.19. Diffraction data were collected on beamlines P11 (Burkhardt et al., 2016) and P13 (Cianci et al., 2017) at PETRA III, DESY (Hamburg, Germany). Data were processed using XDS (Kabsch, 2010). The crystal structures were solved by molecular replacement using the wild-type human P2 structure (PDB entries 3NR3 or 4BVM) (Ruskamo et al., 2014) as a search model in Phaser (McCoy et al., 2007). The structures were refined and built using phenix.refine (Afonine et al., 2012) and Coot (Emsley and Cowtan, 2004). MolProbity (Williams et al., 2018) was used for structure validation.

### Structure of perdeuterated P2 at room temperature

Human P2 perdeuteration, purification, and crystallisation have been described before (Laulumaa et al., 2015a). Crystallisation involved feeding with fresh protein over a period of 10 months to obtain large crystal volumes of 0.3 mm^3^. Neutron data collection and processing have been described (Laulumaa et al., 2015a). X-ray diffraction data were collected from another perdeuterated P2 crystal from the same batch on a GeniX Cu HF rotating anode instrument, and the data were processed with XDS. Joint X-ray/neutron refinement was carried out in phenix.refine, and rebuilding in Coot.

### Small angle X-ray scattering combined with size-exclusion chromatography

SEC-SAXS data were collected on B21 beamline Diamond Light Source (Chilton, Oxfordshire, United Kingdom) (Cowieson et al., 2020). The concentrations of P2-wt, P2-I50del, P2-M114T, and P2-V115A were 9.9, 9.6, 10.5, and 8.9 mg/ml, respectively. The proteins were run in buffer containing 300 mM NaCl, 20 mM HEPES pH 7.5, 1 mM DTT on a Superdex 200 Increase 3.2 column, while SAXS data were continuously collected from the eluate. The data were analyzed with the ATSAS package (Franke et al., 2017) and dummy atom models built with DAMMIN (Svergun, 1999). The data and models were deposited at SASBDB (Kikhney et al., 2020).

### Circular dichroism spectroscopy

For CD measurements, the protein samples were dialyzed with 500-1000 Da Thermo Scientific™ Slide-A-Lyzer™ MINI Dialysis Devices into buffer containing 10 mM sodium phosphate pH 7.7. The proteins were diluted to a final concentration of 50 μg/ml. CD spectra were collected using Hellma quartz cuvettes with a 1.0-mm pathlength between 190-280 nm. For thermal scans, a ramping rate of 1 °C/min between + 22-94 °C was used. Experiments were done with the Chirascan™ CD Spectrometer (Applied Photophysics Ltd). Global 3 (Applied Photophysics) was used to calculate melting temperatures.

### Lipid vesicle aggregation assay

To study vesicle aggregation, a protein concentration series (0, 2.5, 5, 10 and 20 μM) was mixed with 0.5 mM unilamellar vesicles (DMPC:DMPG 1:1), containing a 1:1 molar ratio of DMPC:DMPG, in HBS buffer (150 mM NaCl, 20 mM HEPES pH 7.5). A Tecan Infinite M1000 Pro plate reader was used to measure the absorbance, shaking before each measurement. The temperature was set at + 30 °C and the wavelength at 450 nm. Six measurements every 5 min were done in triplicates. The data were plotted with GraphPad Prism 8.

### DAUDA binding assay

A fluorescent fatty acid analog, DAUDA, was used to study fatty acid binding to P2. The proteins were diluted into HBS. Protein concentrations of 0, 1, 2.5, 5, 10, and 20 μM were used with 10 μM DAUDA. The fluorescence emission spectra (400-700 nm) with excitation at 345 nm for samples were measured with the Tecan Infinite M1000 Pro plate reader. The data were visualised and analysed with GraphPad Prism 8.

### Bioinformatics

Sequences were aligned using ClustalW (Thompson et al., 2002) and ESPript (Gouet et al., 1999). Sequence-based predictions of protein flexibility were done using DynaMine (Cilia et al., 2013).

### Molecular dynamics simulations

The crystal structures of P2-wt and the disease variants, including a bound palmitate molecule, were used as starting points for MD simulations in YASARA, as previously described (Hallin et al., 2021). The AMBER14 force field (Maier et al., 2015) with the explicit TIP3P solvent model was used, and pressure and temperature were controlled with the YASARA densostat (Krieger and Vriend, 2015). The simulated systems were built in a dodecahedral cell, with a physiological ionic strength of 0.15 M NaCl. After energy minimisation, MD simulations were run for at least 1100 ns each, at +25 °C, and further trajectory analysis was done using YASARA. The first 500 snapshots (125 ns) of each simulation were taken as the equilibration period, based on lack of major fluctuations in R_g_ or RMSD values before that point. Thereafter, 1000 ns of the simulation were included in the analysis in all cases.

### Double supported model membrane patches and time-lapse fluorescence microscopy

Hydrated double supported model membrane patches were prepared using the spin coating technique following the method previously described (Berg Klenow et al., 2020; Boye et al., 2018). Briefly, planar mica substrates were glued on glass coverslips and prior to use the mica was freshly cleaved. 30 μL of a 10 mM lipid solution containing 90% DOPC, 10% DOPS, and 0.5% DiD-C_18_ (1,1’-dioctadecyl-3,3,3’,3’-tetramethylindodicarbocyanine, 4-chlorobenzenesulfonate salt) was spin coated at 3000 rpm for 40 s leading to formation of a dry lipid film. After ~12 h in vacuum, the dry lipid film was placed in a liquid chamber and hydrated using a 10 mM Tris buffer (2-amino-2-hydroxymethyl)-propane-1,3-diol), 140 mM NaCl, 2 mM Ca^2+^, pH = 7.4, at +60 °C for 2 h. The hydrated multilayered membrane was gently flushed with buffer to reduce the multilayers of the membrane to the desired structure of double supported membrane patches. Next, the buffer was exchanged ~10 times to remove the excess of floating lipid fragments in the chamber. The samples were then equilibrated at room temperature for 1 h, and the response of the membrane patches to the addition of protein was monitored with time-lapse fluorescence microscopy.

For imaging, a Nikon ECLIPSE TE2000-U inverted fluorescence microscope (Nikon Corporation, Tokyo, Japan) was used. The setup includes a switchable monochromatic Xenon lamp (PolychromeV, Till Photonics GmbH, Grafeling, Germany) for excitation, fitted with a custom filter cube for imaging at 640 nm (DiD). For all experiments, a 40× air objective (Nikon ELWD, NA = 0.60, Plan Fluor and infinity corrected) was used. Images were recorded using a cooled EMCCD camera system (Sensicam em, 1004 × 1002 pixels, PCO-imaging, Kelheim, Germany). The recording was controlled with the associated Live Acquisition software (FEI GmbH, Hillsboro, OR, USA). Time-lapse videos were processed with FIJI (National Institutes of Health, Bethesda, MD, USA).

## Supplementary information

**Table S1. Primers for mutagenesis**

**Movie S1-S4. Time-lapse films of P2 variants interacting with supported double membrane bilayers.**

## Acknowledgements

The use of the facilities and expertise of the Structural biology and Proteomics and protein analysis core facilities, as well as Sequencing Center at the Biocenter Oulu, a member of Biocenter Finland, are gratefully acknowledged. We acknowledge DESY (Hamburg, Germany), a member of the Helmholtz Association HGF, for the provision of experimental facilities. Parts of this research were carried out at PETRA III and we would like to thank beamline staff for assistance in using the P11 beamline. The synchrotron SAXS data were collected on beamline P12 operated by EMBL Hamburg at the PETRA III storage ring (DESY, Hamburg, Germany). We would also like to thank Diamond Light Source (Oxfordshire, United Kingdom) for SAXS beamline B21, and the staff of SAXS beamline B21 for assistance with testing and data collection. Beamtime at ILL is gratefully acknowledged. We acknowledge financial support from the Independent Research Fund Denmark (DFF-FNU), grant no. 7014-00036B (MBK,ACS), Biocenter Oulu (PK,SR,MU), Jane and Aatos Erkko Foundation (PK), and European Spallation Source (PK,SL).

