## Supplementary Table 1 for "Human myelin protein P2: From crystallography to time-lapse membrane imaging and neuropathy-associated variants"

**Supplementary Table S1. Forward primers for the P2 variants for mutagenesis.**

| **Mutation** | **Direction of the primer** | **Sequence** |
| --- | --- | --- |
| I50del | Forward | 5' GCAAGAAAGG AGATATAACT ATACGAACTG AAAG 3’ |
| M114T | Forward | 5' GCTAGTG AATGGGAAAA CGGTAGCGGA ATGTAAAATG 3’ |
| V115A | Forward | 5' CTAGTG AATGGGAAAA TGGCAGCGGA ATGTAAAATG AAG 3’ |
